# MultiStageSearch: a multi-step proteogenomic workflow for taxonomic identification of viral proteome samples adressing database bias

**DOI:** 10.1101/2024.05.15.594287

**Authors:** Julian Pipart, Tanja Holstein, Lennart Martens, Thilo Muth

## Abstract

The recent years, with the global SARS-Cov-2 pandemic, have shown the importance of strain level identification of viral pathogens. While the gold-standard approach for unkown viral sample identification remains genomics, studies have shown the necessity and advantages of orthogonal experimental approaches such as proteomics, based on proteomic database search methods. The databases required as references for both proteins and genome sequences are known to be biased towards certain taxa, such as pathogenic strains or species, or common model organisms. Aditionally, the proteomic databases are not as comprehensive as the genomic databases.

We present MultiStageSearch, an iterative database search approach for the taxonomic identification of viral samples combining proteomic and genomic databases. The potentially present species and strains are inferred using a generalist proteomic reference database. MultiStageSearch then automatically creates a proteogenomic database. This database is further pre-processed byfiltering for duplicates as well as clustering of identical ORFs to address potential bias present in the genomic database. Furthermore, the workflow is independent of the strain level NCBI taxonomy, enabling the inference of strains that are not present in the NCBI taxonomy.

We performed a benchmark on several viral samples to demonstrate the performance of the strain level taxonomic inference. The benchmark shows superior performance compared to state of the art methods for untargeted strain level inference using proteomic data while being independent of the NCBI taxonomy at strain level.

## Introduction

Viral and bacterial pathogens represent one of the biggest threats to public health,^1^ as evidenced by the recent SARS-CoV-2 pandemic. In suspected cases of outbreaks, the availability of fast and accurate detection and identification methods for these pathogens is essential.^2^ Routinely used diagnostic methods such as Rt-PCR^3^ have the disadvantage of requiring previous knowledge of the sought-out pathogens. ^4^ Therefore, especially for applications in viral surveillance and the detection of novel or uncommon pathogens, open-view approaches such as genomics and proteomics come into play. ^5^ While, especially for viral surveillance, genomics has established itself as gold-standard method,^6^ recent studies have demonstrated the advantages of proteomics as an orthogonal approach. ^7,8^

With continuous advances on the instrumentation level, tandem mass-spectrometry preceded by liquid chromatography (LC-MS/MS) is the current method of choice for proteomic analysis.^9^ It is high-throughput and offers high resolutions at the peptide and therefore protein sequence levels. LC-MS/MS has been successfully used to identify viral pathogens, both at MS1^10–12^ and MS2 levels.^13^

The standard downstream bioinformatic analysis of proteomic samples relies on a search of the MS/MS spectra against proteomic reference databases.^14^ Different versions of comprehensive reference databases exist, and many are available online. For databases like NCBI^15^ and Uniprot,^16^ there are curated subsets, where the proteomic references and their taxonomic assignments have passed a quality control.^17,18^ However, the curated subsets can have a limited range regarding their representation of protein sequence and taxonomic diversity, especially when researching organisms less common for proteomic applications, as is the case for viral or metaproteomics.^19–21^ It can therefore be beneficial to include all, also the uncurated, available proteomes for taxonomic identifications.^8,22^

To leverage all the proteome information available, workflows using a two-step search approach have been developed.^23^ Usually, they use a first proteomic database search against a database with less taxonomic resolution but a broad representation across possibly present species,to identify potentially present taxa (or proteins).^22^ This is followed by a second search against all higher-resolved proteomes available for appropriate candidate species. While this strategy has shown to be successful for many samples, it can lead to wrong taxonomic attributions. Previous studies^8^ have suggested that this may be the results of bias present in the reference databases. In fact, the presence of bias towards model organisms in proteomic and genomic databases has been previously described.^24^ Any researcher can upload proteomes to UniprotKB or NCBI, a practice that is encouraged since it will lead to a broad representation of proteomic diversity and widen taxonomic diversity by including more species, strains and isolates. However, this practice introduces bias. Strains or species towards which databases are biased can be, for example, ones that are easily available for ordering through certain distributors or ones that very commonly infect people,^25^ such as certain Influenza strains.^26^ The overabundant presence of reference proteins for a certain strain will increase the probability of finding spectra associated to it and bias the taxonomic analysis results. These reference-related biases have been described for genomics^27,28^ and are equally valid for proteome reference databases.

This bias is known, and possible solutions proposed include the clustering of overly similar references to reduce over representation.^29^ Another potential avenue to address this database bias and leverage additional references available is to use genomic databases. While these may be biased as well, genomics is more established for the analysis of viral samples, leading to a more comprehensive representation of viral protein, strain and species diversity.^30^

To make use of additional genomic information and manage bias present in proteomic databases, we introduce MultiStageSearch. MultiStageSearch is a multi-step proteogenomic workflow for strain level-identification of viral samples. In a first step, a traditional proteomic database search is performed against a generalist proteomic reference database. Based on the resulting matches, potential candidate taxa are identified and a proteogenomic database is constructed using six-frame translation. Tailored strategies are used to identify, filter and download fitting reference genomes for database construction. This results in taxonomic identifications for single organism samples that have the potential to overcome representation bias in proteomic, but also genomic databases.

MultiStageSEarch is written in python and implemented as a snakemake workflow. The code is available at https://github.com/rki-mf2/MultiStageSearch

## Methods

### Workflow Overview

We start by introducing the MultiStageSearch workflow, providing a detailed description of every search step.

MultiStageSearch performs a total of up to four database search steps to yield strain level identification results for experimental MS spectra. During the workflow, strain level genomes are automatically queried and downloaded from the NCBI nucleotide database^31^ with two different approaches to perform a proteogenomic database search.

The input to MultiStageSearch is a MGF file containing the MS spectra, a parameter file for SearchGUI, a reference database (for example RefSeqViral^32^) as well as a file that maps the protein accessions to the Taxon-IDs and optionally a host and CRAP^33^ database. Each search step (Host Filtering, Reference DB Search, Genomic DB Search, Top-Scoring DB Search) uses SearchGUI^34^ to add decoys to the provided database for the target-decoys search approach. ^35^ Afterwards SearchGUI is called using X!Tandem^36^ to perform the database search. Post-processing is either performed by PeptideShaker^37^ or MS2Rescore.^38^ While MS2Rescore requires additional configuration for each sample such as the regular expression to match the spectrum titles, when choosing PeptideShaker, MultiStageSearch is able to perform the database search for multiple samples at once.

The Workflow is build in Snakemake^39^ and can be broadly divided into the steps shown in Figure 1. The code is written in python and tested for python version 3.9. The computations were performed on the linux compute servers of the Freie Universität Berlin. Additional packages used in this workflow are summarized in table 3 in the supplementary materials. The detailed workflow steps are described in the following.

**Figure 1:**
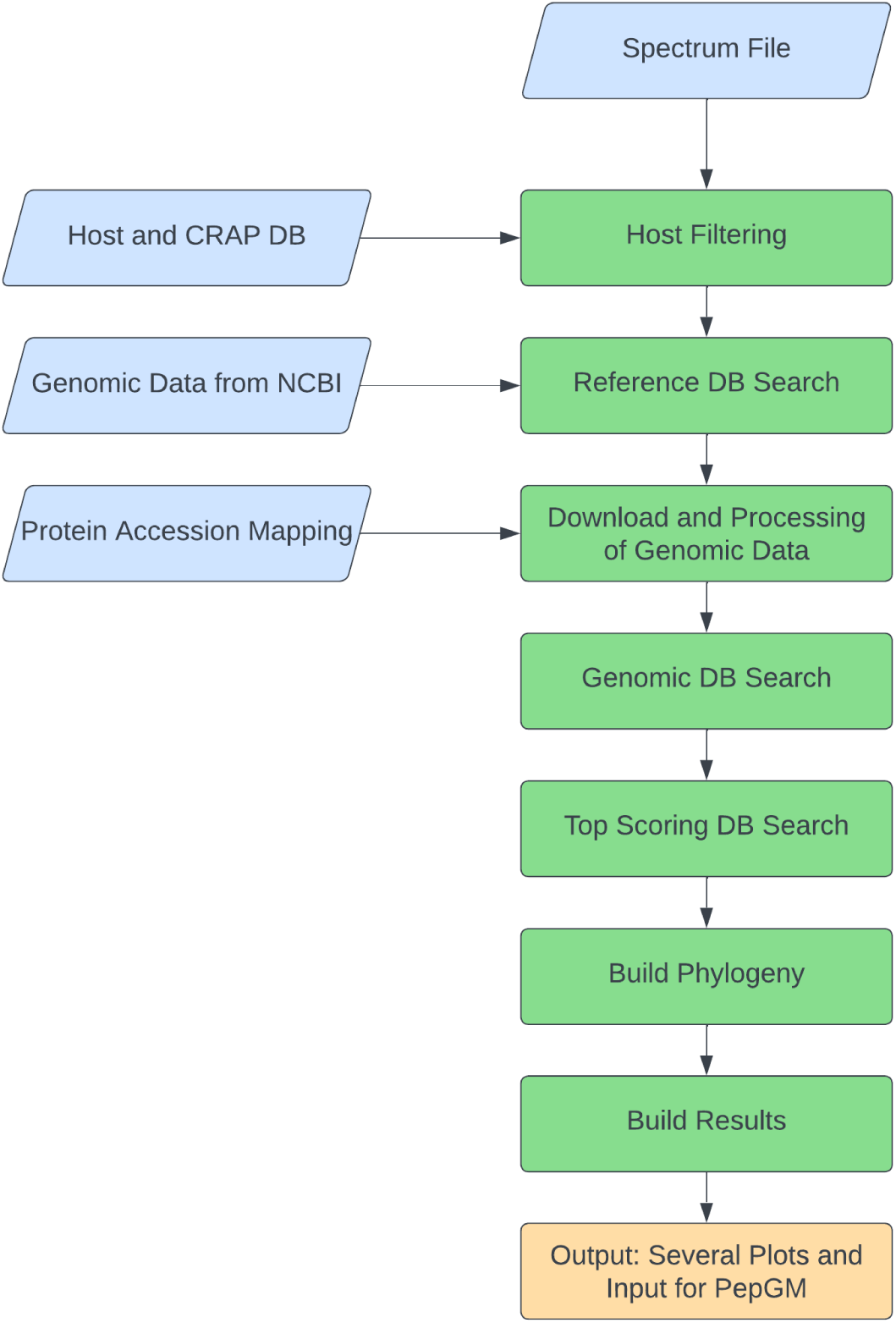
Overview of the MultiStageSearch workflow. In blue, the input to the workflow is shown. Green boxes correspond to the main steps of the workflow. In Orange, the output is summarized.

### Host Filtering

The user can choose to filter the experimental spectra, input in .mgf format, for a user-chosen host and contaminant database. After searching and identifying Peptide-Spectrum-Matches (PSMs), those PSMs are filtered out from the original MGF file. This Host Filtering is optional and is turned on by default. If the Host Filtering is disabled, the Reference DB Search is performed on the original MGF file.

### Reference DB Search

Using the user-provided .fasta reference database, MutliStageSearch performs the database search as described above. Here, it is recommended to use a a reference database spanning a broad range of viral species such as RefSeqViral for viruses. If there is prior knowledge about the sample, e.g. the host species or potentially present host symptoms, this database can be reduced accordingly. PSMs are imported into MS2Rescore or PeptideShaker, which are used for post-processing, rescoring and results export.

The resulting report of PeptideShaker or MS2Rescore contains a list of PSMs. From these PSMs, appropriate candidate species for the further workflow execution need to be inferred. To this end, MultiStageSearch uses a weighted aggregation of PSM counts per taxon, similarly to^40^ and.^8^ The weight of a PSM is based on it’s degeneracy, meaning the more proteins a PSMs can be attributed to, the less weight it will have. Afterwards they are mapped to the corresponding species Taxon-IDs. To this end, the proteins should contain the NCBI identifiers, as is the case for RefSeqViral. The weights for each Taxon-ID are summed and written to a tsv file, which builds the input for the next step. The user-defined threshold ‘max weight differences’ determines how many taxa will be used to identify candidate strains for the next analysis steps. Since, in many cases, the weight differences between the Taxon-IDs are high, this threshold ensures that MultiStageSearch considers as few Taxon-IDs as required. The indentified candidate taxa are used as input for the next workflow step.

### Download and Processing of Genomic Data

#### Download of candidate strain genomes

In this step, strain-level genomes for the top-scoring species-level Taxon-IDs are automatically queried from the NCBI database. Several challenges present themselves when downloading the appropriate genome sequences for the previously identified candidate taxa. All genome references available in NCBI are taken into consideration. For each, a tailored solution to increase the robustness and replicability of the workflow were developed. These solutions are presented in the following. Several user-adjustable parameters define the query.

- number of taxids: defines the maximum number of species taxa to take into account for the query.
- max weight differences: defines the maximum weight difference between the last Taxon-ID considered by the workflow and the next top-scoring Taxon-ID. If the difference is to great, the next Taxon-ID will not be considered.
- max number accessions: defines the maximum number of genbank^41^ accession that will be processed.
- sequence length diff: defines the maximum sequence length difference between the last sequence considered by the workflow and the next longest sequence. If the difference is to great, the next sequence will not be considered.
- max sequence length: defines the maximum sequence length. This can be used to avoid whole genome shotgun sequence entries.
- use NCBI Taxa: Use the NCBI Taxonomy^15^ for the query.
- only use complete genomes: search only for sequences with “complete” in the title (e.g. “complete genome”)

In the NCBI database, genomes are often not linked correctly to the corresponding Taxon-ID of the strain, but to the species. Therefore, MultiStageSearch uses the title of the database entry instead of the Taxon-ID. If use NCBI Taxa is set to “True”, MultiStageSearch first finds all (alternative) names for the species. Afterwards, all combinations of species name and corresponding strain name from the NCBI taxonomy are queried and the sequences matching the parameters are downloaded. If use NCBI Taxa is set to “False”, MultiStageSearch will search for sequence entries with the Taxon-ID of the species. This approach is able to find more entries and also strains that are not present in the NCBI taxonomy.

Using different information from the database entries, a tsv file containing the values gen-bank accession, species, strain, isolate, taxa and HigherTaxa, for each downloaded genome, is generated. Taxa and HigherTaxa correspond to the strain Taxon-ID and species Taxon-ID. While the strain Taxon-ID can be inferred when using the “use NCBI Taxa” option, there are no Taxon-IDs available for most strains when the “use NCBI Taxa” option is set to “false”. Therefore, in this case, own Taxon-IDs are created. MultiStageSearch assigns a Taxon-ID, starting at 5.000.000 (to avoid overlap with existing Taxon-IDs), to each downloaded sequence.

#### Generation of the proteogenomic reference database

Since proteome reference databases are required as input for the second database search step, Sixpack^42^ is used to perform a six-frame translation for converting the nucleotide sequences to proteomes. These are then concatenated for further processing.

The dowload approach describes above often results in multiple genomes, and therefore proteomes, for the same strain. To avoid these duplicate proteomes after the six-frame translation, a filtering is implemented. The strain and isolate names from the tsv file generated during the genome download are compared. If they are identical, the proteomes are compared. To this end, every open reading frame (ORF) of proteome 1 is compared to every ORF of proteome 2. If the sequences of the ORFs are identical, this pair increases a counter *c* by one. In the end, the pairwise similarity *S_P1, P2_* of the proteomes is computed by 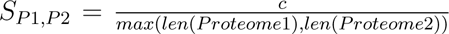, where the length corresponds to the number of ORFs in the proteomes. If the user defined similarity threshold (default 98%) is exceeded, only the longer proteome of the identical proteomes is used for further analysis. For the obtained proteome database, SearchGUI is used to create the decoys. The resulting database still contains a great number of identical ORFs, since the downloaded genomes correspond to strains with the same parent species. To reduce the database size and address the high number of duplicate ORFs, these ORFs are clsutered. If ORFs have the same sequence, the sequence headers are combined to retain the information, and the sequence is stored only once.

To be able, to map the ORFs to the strains again, a mapping file is created, analogue to the approach for the reference database search.

### Genomic DB Search

The downloaded and processed strain genomes are now the basis of the next database search. Since genomics is more established than proteomics,^43^ typically, more genomes will be found at strain level using this proteogenomic approach, increasing the number of PSMs. The resulting PSMs are also weighted and aggregated by Taxon-ID using the same weighting as the first search step. This results in a list of top-scoring, strain-level taxa.

### Build Phylogeny

Optionally, a phylogeny will be build upon the *n* (default=30) top-scoring strain taxa. Therefore, the strain genome fasta file is filtered for the top-scoring strain taxa. Afterwards, MAFFT^44^ is called in auto mode to perform a multiple sequence alignment (msa). This msa is then used by IQ-TREE^45^ to reconstruct the phylogenetic tree on strain level. While the nodes of the phylogentic tree are now named after the genbank accessions of the corresponding genomes, they are renamed to match the corresponding Taxon-IDs. This can be important for further downstream tools, e.g. PepGM.

### Top-Scoring DB Search

Since the phylogeny is only built upon the *n* top-scoring strain taxa, a lastsearch step is performed optionally. This search step only uses the proteomes of the top-scoring strain taxa from the second search. Otherwise it performs the same steps as the genomic database search step, resulting in a list of the *n* candidate strains with corresponding weighted PSM scores. Additionally to the reasoning of the phylogeny, it can enhance the results, since the database now contains fewer data and fewer duplicates. This leads to more PSMs for the taxa present in this new proteogenomic database, potentially providing more reliable results. The taxon with the highest number of weighted PSMs is then identified as present.

### Database Suitability

The user might not always be sure of the quality of the sample and whether the correct databases are provided as input. To verify their quality, MultiStageSearch is able to compute a modified version of the protein sequence database suitability described in .^46^ To this end, NOVOR^47^ is used to compute *de novo* peptides from the provided mgf file containing the spectra. These de novo peptides are given a unique header containing *”NOVOR PEPTIDE “* as a prefix. they are then concatenated with the database used as input for the current search step. Afterwards, the standard database search steps are applied to this modified database. Using the resulting report of PeptideShaker or MS2Rescore, the proportion of “real” database hits and NOVOR peptide hits is computed.

While^46^ performs a re-ranking of PSMs with almost identical scores, this is omitted in MultiStageSearch. To show the suitabilities of each database used, the computation of database suitability is performed at each search step. The calculated database suitability value can indicate to the user if either the reference database used is inappropriate, or the spectra present in the mgf file are of very poor quality.

### Results output and visualization

Finally, to visualize the results of the database searches, the similarity comparisons and the suitability of the databases, several plots are created. These show the different results: the weighting of taxa for the different search steps, the similarities of proteomes and identified peptidomes, and other metrics. The similarty of the identified peptidomes is calculated similarly to the pairwise similarity of the proteomes *S_P_* _1_*_,P_* _2_, except only identified ORFs are considered.

Additionally to the plots, MultiStageSearch creates output that is readable by potential further downstream analysis pipelines such as PepGM.^8^ The input files for PepGM are the phylogenetic tree in newick format, the csv file containing the accessions and Taxon-IDs for the strains, a csv file containing the final scores (weights) for the top-scoring strains as well as a csv file containing the identified peptides, the sequences of the peptides, the score of the PSMs, the Taxon-IDs of the strain and the ancestor.

### Benchmarking Samples

To benchmark the performance of MultiStageSearch against other tools like TaxIt ^22^ and PepGM, several viral samples were tested. Among those were two Cowpox samples (PXD003013, PXD014913) of which both are the strain “Brighton Red”, a Hendra virus sample, (PXD001165, strain “HeV/Australia/1994/Horse18”), an avian bronchitis sample (PXD002936, strain “Beaudette CK”), a Adenovirus sample (PXD004095, strain “2”), a Herpes simplex 1 sample (PXD005104, strain “F”) and two SARS-CoV-2 samples (PXD018594, PXD025130) both of lineage B. All samples can be found in the PRIDE^48^ database.

## Results

To showcase the MultiStageSearch taxonomic identification abilities, we benchmarked MultiStageSearch against PepGM and TaxIt. To this end, we use the 8 samples as described in section “Benchmarking Samples”. While MultiStageSearch is able to identify the correct strains of most samples as shown in table 1, TaxIt was not benchmarked on the SARS-CoV-2 samples. PepGM as well as MultiStageSearch were only able to identify the correct species.

**Table 1:** Identification results of the different methods.

| Sample | MSS with Taxonomy | MSS without Taxonomy | TaxIt | PepGM |
| --- | --- | --- | --- | --- |
| PXD003013_Cowpox | ST+ | ST+ | ST+ | ST+ |
| PXD014913_Cowpox | ST+ | ST+ | ST+ | ST+ |
| PXD001165_Hendra | ST+ | ST+ | ST+ | ST+ |
| PXD002936_Avian | SP+ST- | SP+ST- | SP+ST- | SP+ST- |
| PXD004095_Adeno | SP+ST- | ST+ | ST+ | ST+ |
| PXD005104_Herpes | ST+ | ST+ | SP+ST- | SP+ST- |
| PXD018594_Covid | SP+ | SP+ | — | SP+ |
| PXD025130_Covid | SP+ | SP+ | — | SP+ |

As shown in table 1, the strain of the avian bronchitis sample could not be identified by any of the methods. Overall, the results show that MultiStageSearch is not dependent on the NCBI Taxonomy and still outperforms TaxIt and PepGM. Notably, MultiStageSearch is able to correctly identify the strain of the Herpesvirus sample, which both TaxIt and PepGM were not. The results show that MultiStageSearch is able to leverage the advantages of genomic data as well as the independency of the NCBI taxonomy to reach higher strain level identification accuracy for most samples benchmarked. The results are discussed in more detail in the following.

### Taxonomic identification using the NCBI taxonomy

When using the NCBI Taxonomy for querying the genomic strain level data in the NCBI nucleotide database, MultiStageSearch is able to identify the correct strain for the two Cowpox samples, the Hendra sample as well as the Herpes sample as shown in Table 1. In Figure 2, the final results for the Herpes sample are shown. Using the weighting described above, the strain “F” is identified, which is the correct strain of this sample. This showcases how the approach used by MultiStageSearch is able to overcome database bias: PepGM identified the herpesvirus strain 17^8^ as present for this sample and discusses how this misidentification is due to database bias. In fact, the herpesvirus 1 strain 17 is distributed by a scientific retailer.^49^ Because it is readily available, it is more readily used in experiments, and corresponding protein sequence are uploaded to online databases in greater number: strain 17 has 670 protein entries in the NCBI database to date (may 2024), but strain F has only 25. The SARS-CoV-2 samples on the other hand cannot be identified on subspecies level using the taxonomy. This is due to the missing data for SARS-CoV-2 in the nCBI taxonomy. While taxonomic data is available for the avian bronchitis and the adenovirus sample, both are not identified correctly. As Figure 3 shows, the two strains “Beaudette CK” and “Beaudette” have the same weight for avian bronchitis. Figure 4 shows the pairwise peptidome similarity between the proteogenomic sequences for the avian bronchitis sample. It is visible that the strains “Beaudette CK” and “Beaudette” have 100% similarity for the identified peptidomes. Therefore, MultiStageSearch as well as TaxIt and PepGM are not able to identify the correct strain “Beaudette CK”.

**Figure 2:**
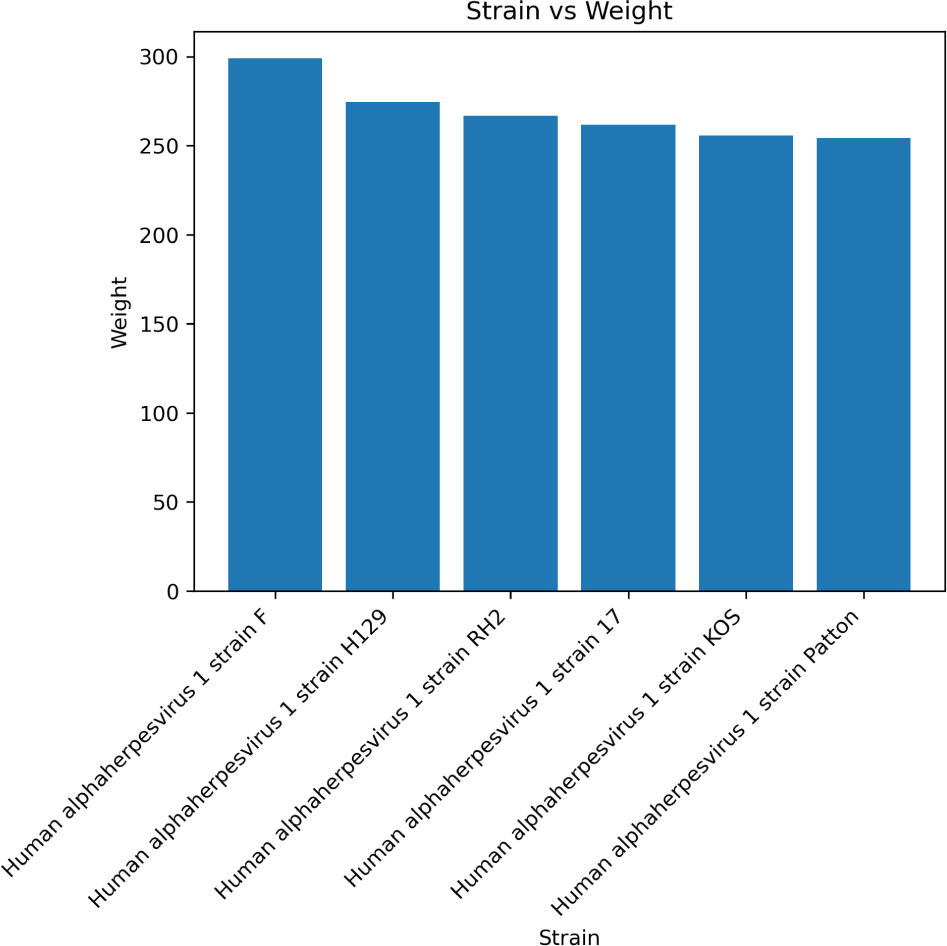
Bar plot showing the final weights of the top-scoring database search for the herpes sample.

**Figure 3:**
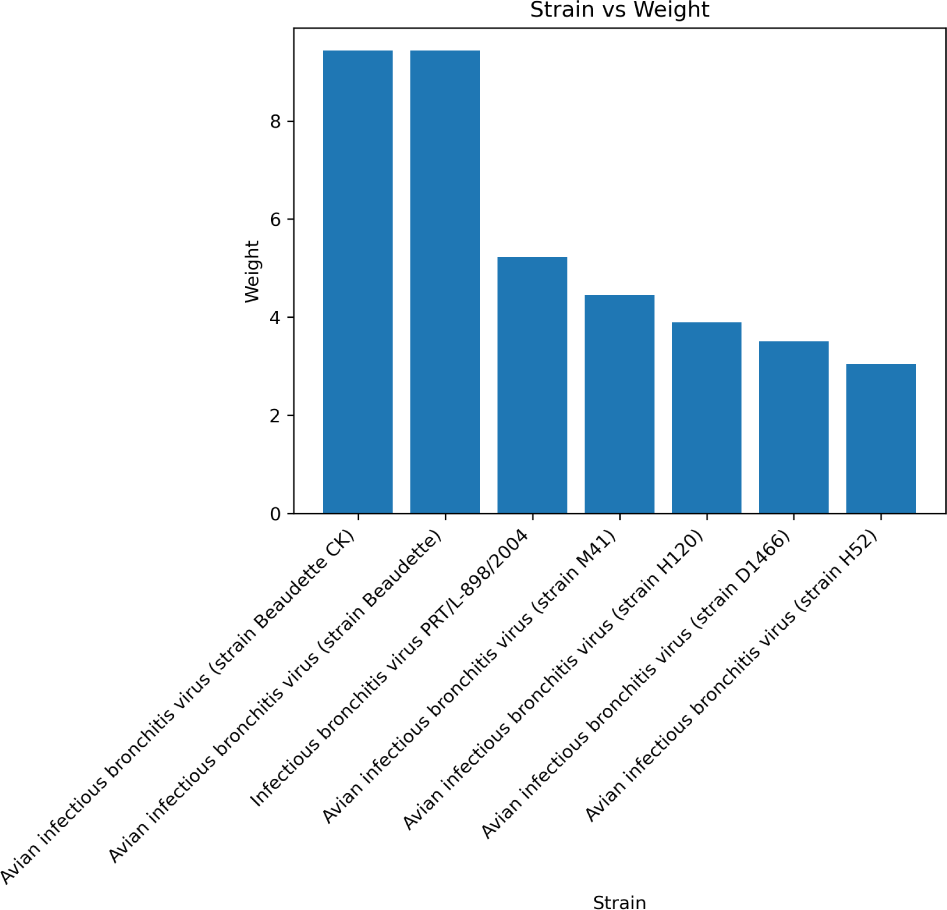
Bar plot showing the final weights of the top-scoring database search for the avian bronchitis sample.

**Figure 4:**
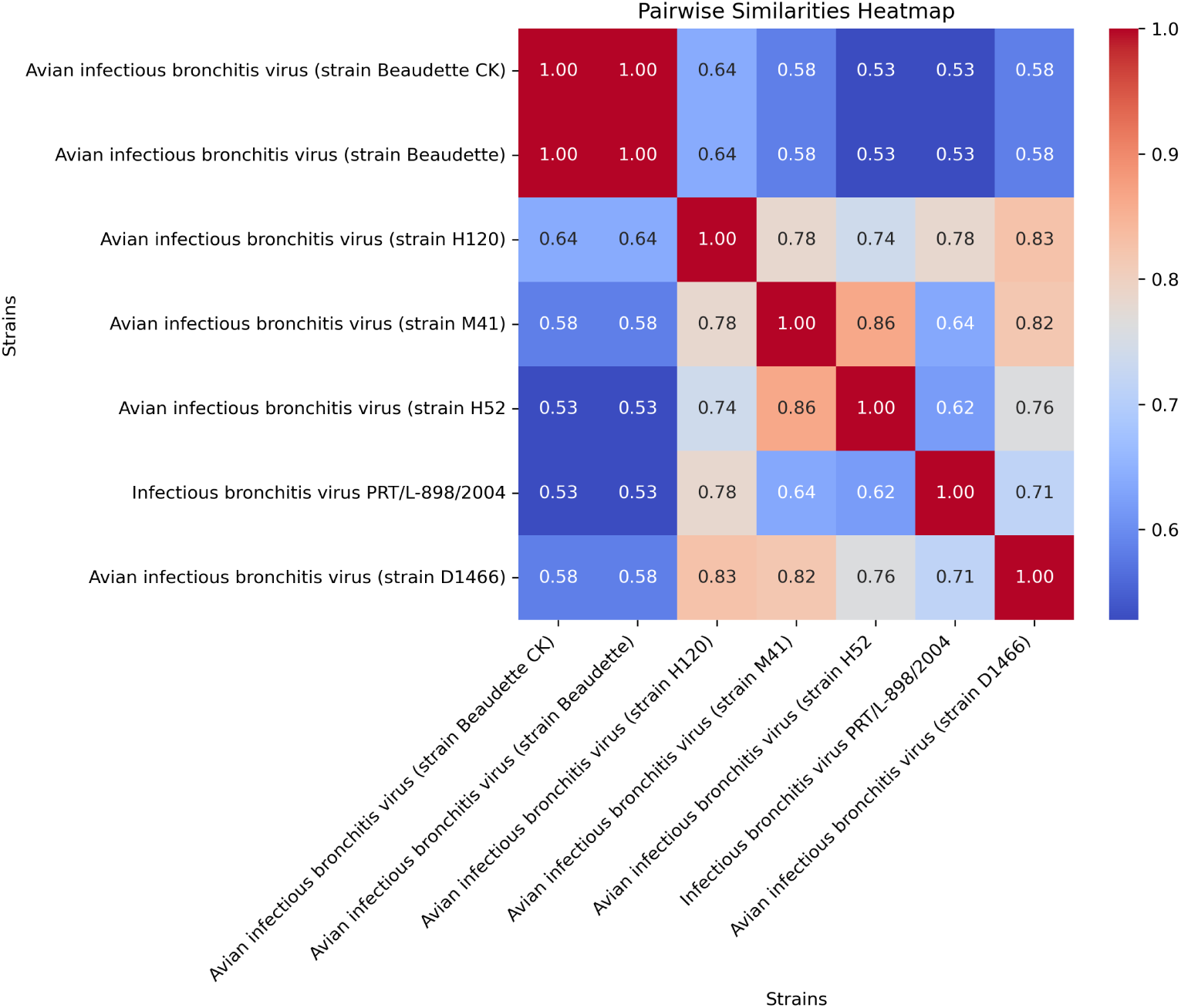
Heatmap of the similarities of identified Peptidomes of the top-scoring genomes for the avian bronchitis sample.

The situation is different for the adenovirus. In this case, the reference database search finds two species with highest weights, as shown in Figure 5. These correspond to “Human mastadenovirus C” and “Human adenovirus 2”. Important to note is that in the NCBI taxonomy, “Human adenovirus 2” is a descendant of “Human mastadenovirus C” without a specified rank. When querying the database using the name combinations as described in section “Download and Processing of Genomic Data”, there are no strain level genomes found, since this query searches for the same names for species and strain. An example for this is:

**Figure 5:**
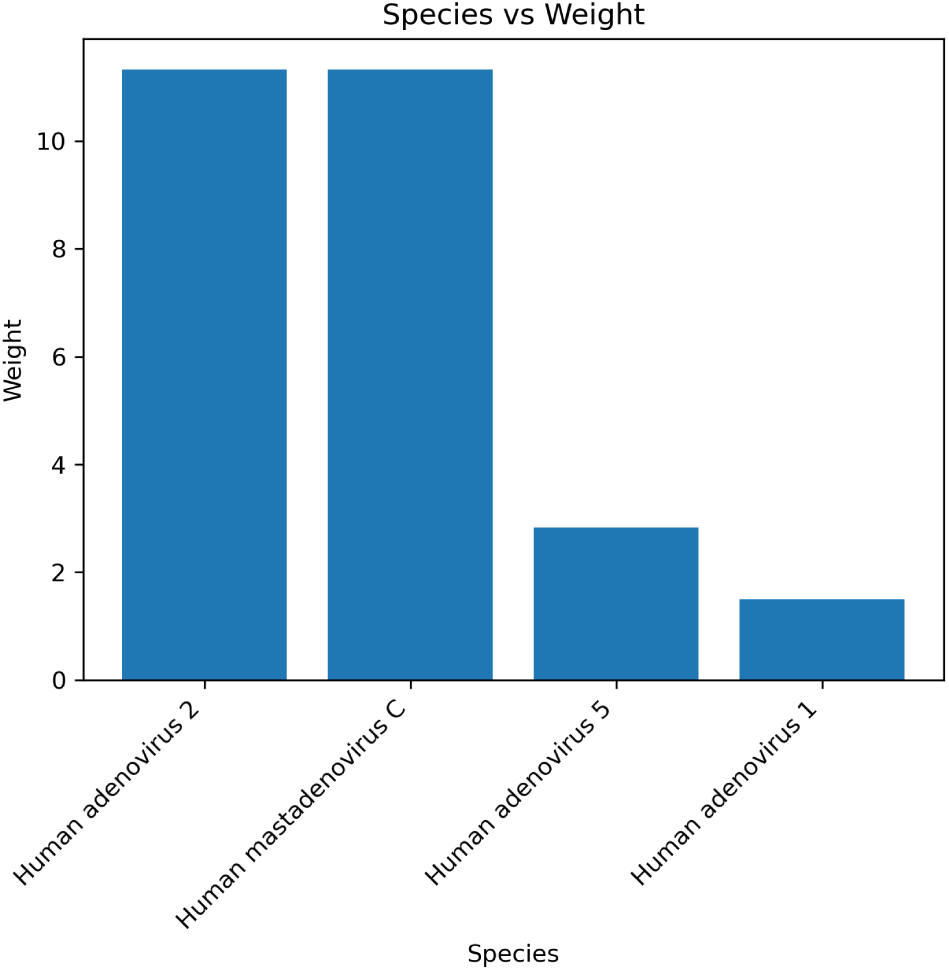
Barplot showing the identification results as species level against the reference database for the adenovirus 2 sample.

Human adenovirus 2 AND ((strain Human adenovirus 2C[All Fields]) or (Human adenovirus 2C[Title])) AND “complete “[Title]

This query strategy therefore does not find the correct strain level genome for the adenovirus sample.

### Taxonomic identification without using the NCBI taxonomy

The results of Table 1 show that MultiStageSearch still finds the correct strain with even higher accuracy than TaxIt and PepGM without the usage of the NCBI taxonomy. Instead of using the strain-level NCBI taxonomy, which is most often inexistant, it searches for entries that are linked to the Taxon-ID of the species. A significantly higher number of strains and isolates were found in the NCBI database using this approach and the strains were still identified correctly for most samples. While the NCBI taxonomy for SARS-CoV-2 Virus at levels lower than species is not defined at the time of writing, SARS-CoV-2 has millions of proteins and nucleotide sequences in the NCBI database. Therefore, the standard query of MultiStageSearch is not able to query the database in a reasonable timeframe and no correct strain can be found using the approach with name combinations of the NCBI Taxonomy. The other virus sample where the virus could not be detected correctly on strain level is the avian bronchitis sample. In this case, a similar problem as for the approach using the NCBI taxonomy occurs. Figure 6 shows two key findings: (1) MultiStageSearch finds only very few PSMs for the top-scoring genomes with only up to 36 PSMs per strain/isolate, likely due to the quality of the sample itself. (2) The three top-scoring strains have the exact same amount of PSMs. As for the approach using the NCBI taxonomy, the identified peptidomes of the top-scoring genomes are identical. Therefore they cannot be distinguished. Furthermore, the filtering of duplicate proteomes does not work here, since the strain names are too different.

**Figure 6:**
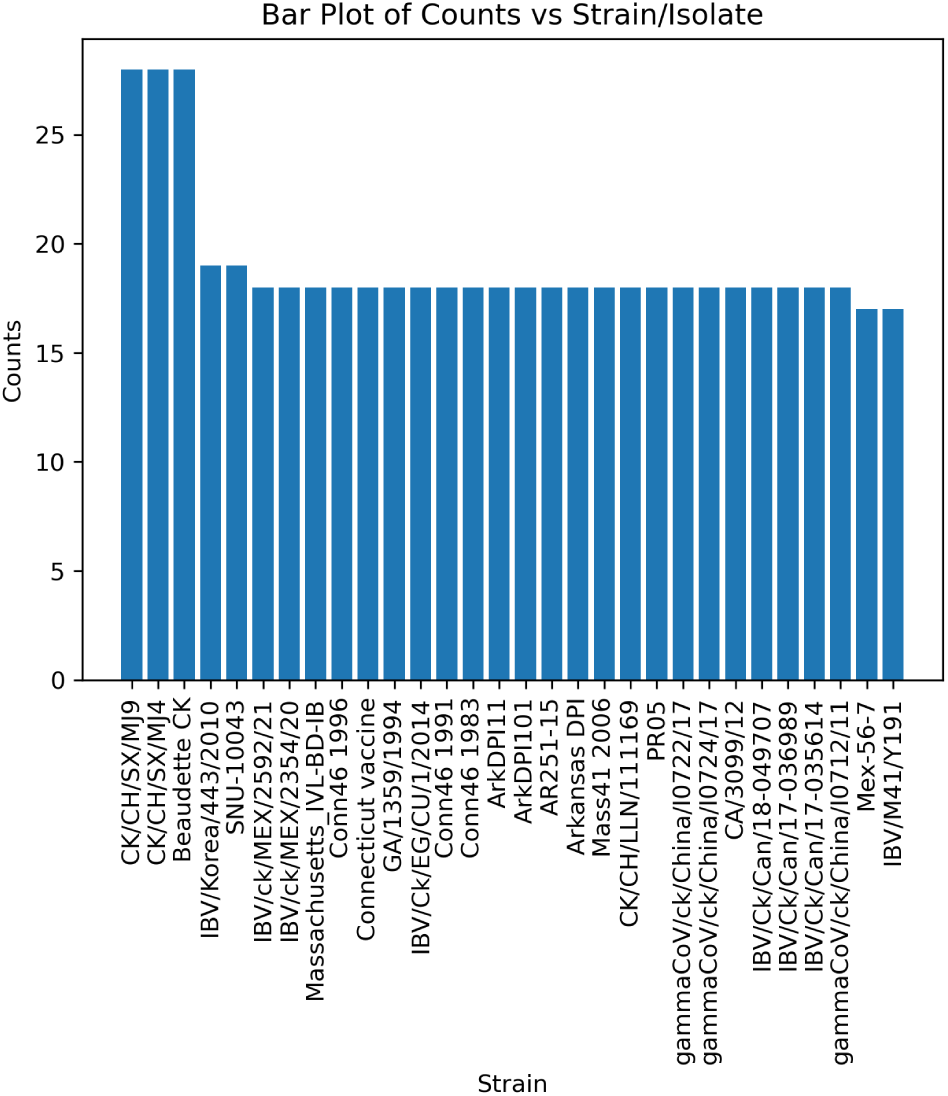
Bar plot showing the number of PSMs per strain/isolate of avian bronchitis.

Table 2 shows the advantages of the combination of the independence of the NCBI taxonomy and the proteogenomic approach. While most viruses do not have many strain references in the NCBI taxonomy,^15^ many more genomes are found that can be assigned to different strains and isolates not represented in the taxonomy. Therefore, the genetic variance of this species is covered to a greater degree and the automatically created proteogenomic database covers more variance for this species. The Cowpox virus has five strains in the NCBI taxonomy and 98 complete Genomes are found using MultiStageSearch. Although there is only one strain (”Hendra virus horse/Australia/Hendra/1994”) present in the NCBI taxonomy for the Hendravirus, MutliStageSearch is able to find 19 complete Genomes. While for the three samples avian bronchitis, adenovirus and herpesvirus hundreds of complete genomes are found, MultiStageSearch finds millions of complete genomes for SARS-CoV-2, showcasing the need for further bioinformatic solutions, beyond the one presented in this work, able to leverage this wealth of information.

**Table 2:** Table of available strains in the NCBI taxonomy as well as the number of found genomes for this species and the number of used genomes.

| Sample | Strains in Taxonomy | Found (complete) Genomes |
| --- | --- | --- |
| PXD003013_Cowpox | 5 | 98 |
| PXD014913_Cowpox | 5 | 98 |
| PXD001165_Hendra | 1 | 19 |
| PXD002936_Avian | 34 | 500 |
| PXD004095_Adeno | 44 | 361 |
| PXD005104_Herpes | 17 | 164 |
| PXD018594_Covid | 0 | 5.364.560 |
| PXD025130_Covid | 0 | 5.364.560 |

**Table 3:**
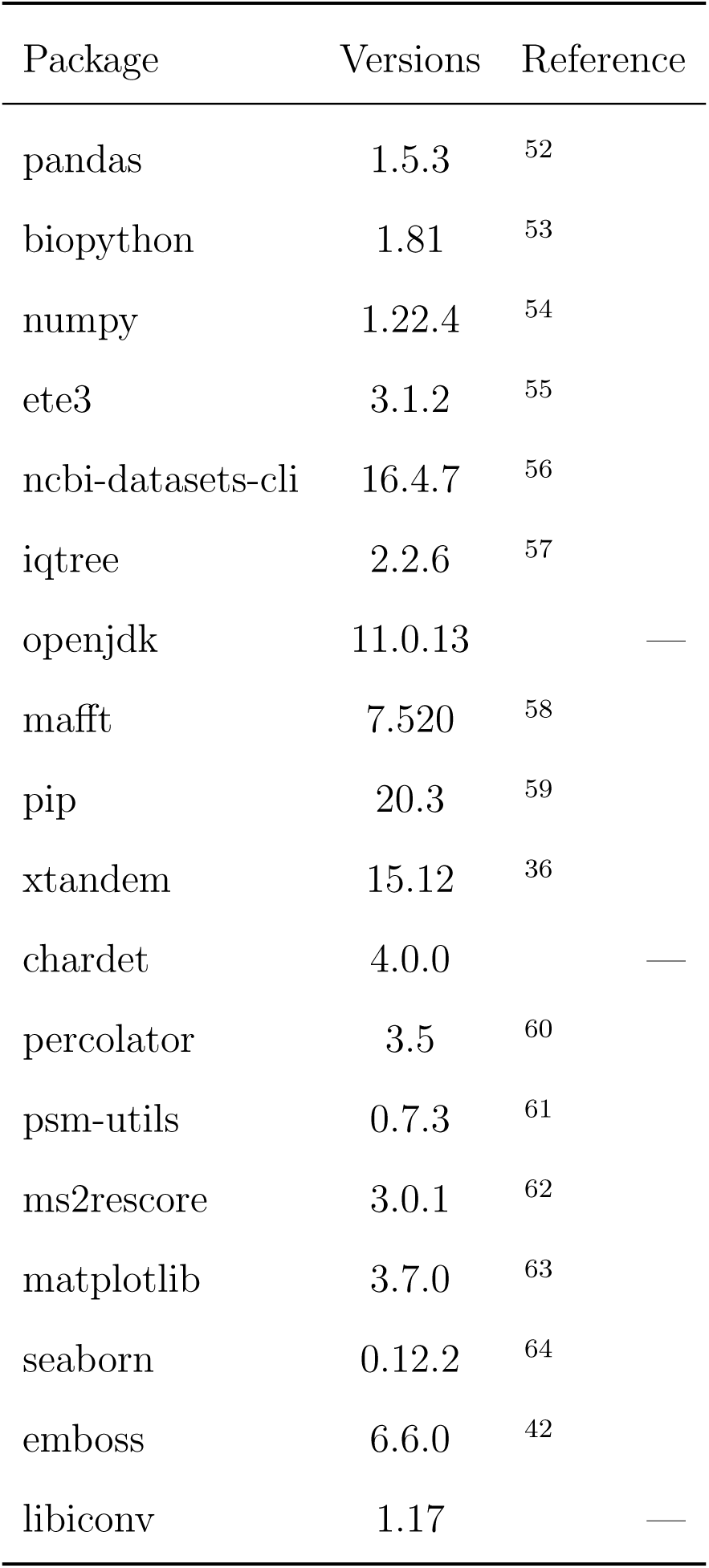

### Database Suitability

The choice of the database influences the identification results. While the user provides the databases for the host filtering and the reference database search, MultiStageSearch creates the genomic database automatically based on the intermediate results. Figure 7 shows the database suitabilities of the four search steps for the PXD003013 Cowpox sample using the query approach without the NCBI taxonomy. The suitabilities were computed following the procedure described in the methods section above. The databases provided by the user, in this case the host (*Bos taurus*) and cRAP database and the RefSeq viral database, have a suitability of 43.85% and 71.28% respectively. The automatically created proteogenomic database has the highest suitability with 93.74%, underscoring the advantage of the proteogenomic database generated.

**Figure 7:**
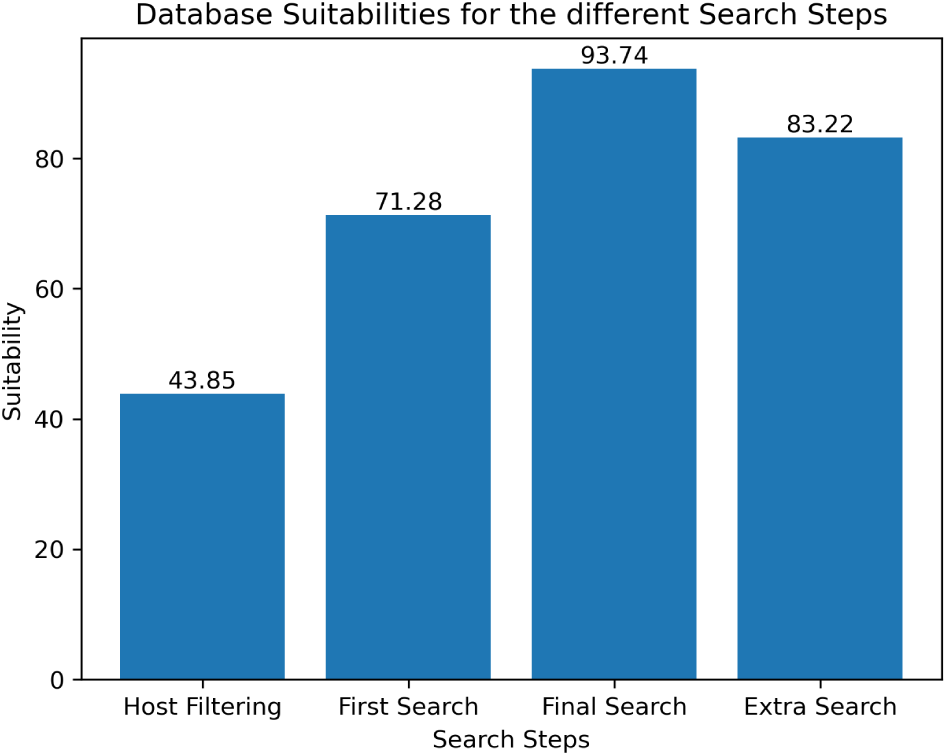
Bar plot showing the database suitabilities of the PXD003013 Cowpox sample using the query approach without the taxonomy.

Figure 8 shows the database suitabilities of the herpes sample. Here it is visible, that the database suitabilities greatly differ. While the suitability of the host database is higher compared to the one for the PXD003013 Cowpox sample in figure 7, the suitability of the reference database is much lower. This can have multiple reasons: (1) The database does not contain the correct species of the sample. (2) The sample contains more spectra that are not from the species analyzed. (3) The overall quality of the sample is low. All possibilities cause a lower database suitability as fewer database peptides are found and the number of identified de novo peptides rises.

**Figure 8:**
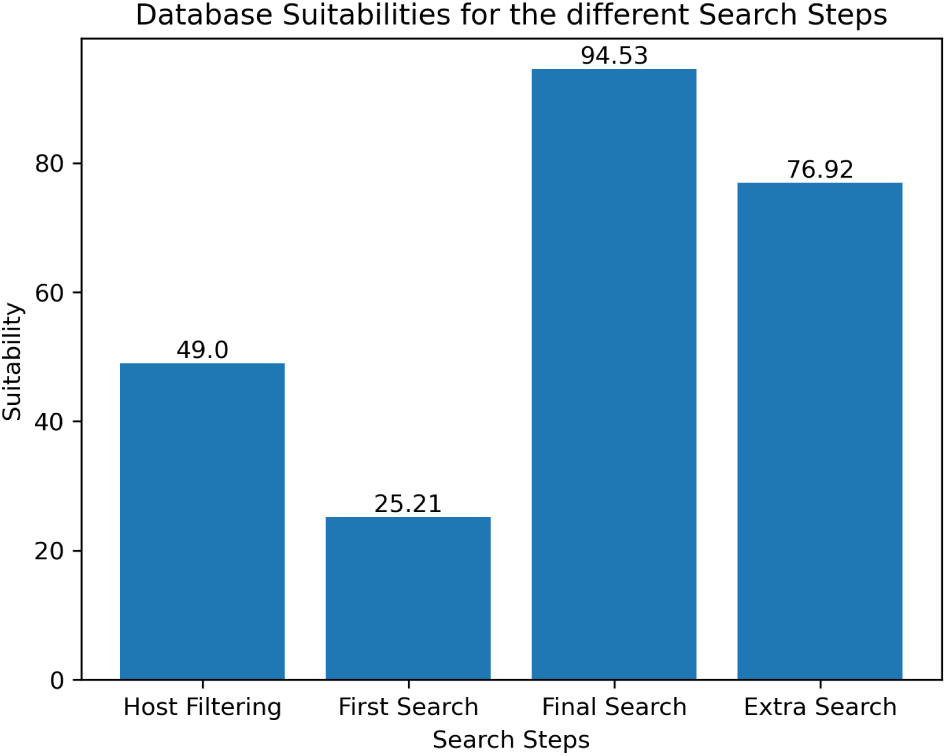
Bar plot showing the database suitabilities of the herpes sample using the query approach without the taxonomy.

On the other hand, the high suitability of 94% of the genomic search shows that it can lead to a more sample appropriate database even when the suitability of the reference database or the quality of the sample are not very high.

Figure 9 shows the database suitabilities for the the hendravirus sample. Here every suitability except the suitability of the host database (41.56%) is very low with only 5.51 to 6.66 percent. This indicates that the sample contains only very little of the virus itself, since the database suitability for the host database is significantly higher compared to the databases containing viral references. Nevertheless, MultiStageSearch is able to find the correct strain “HeV/Australia/1994/Horse18”, which might partly be due to the relatively low number of strains available as a reference additionally to the correct reference strain (19 compared to 98 for the next lowest number of reference for the cowpox virus, refer to table 2).

**Figure 9:**
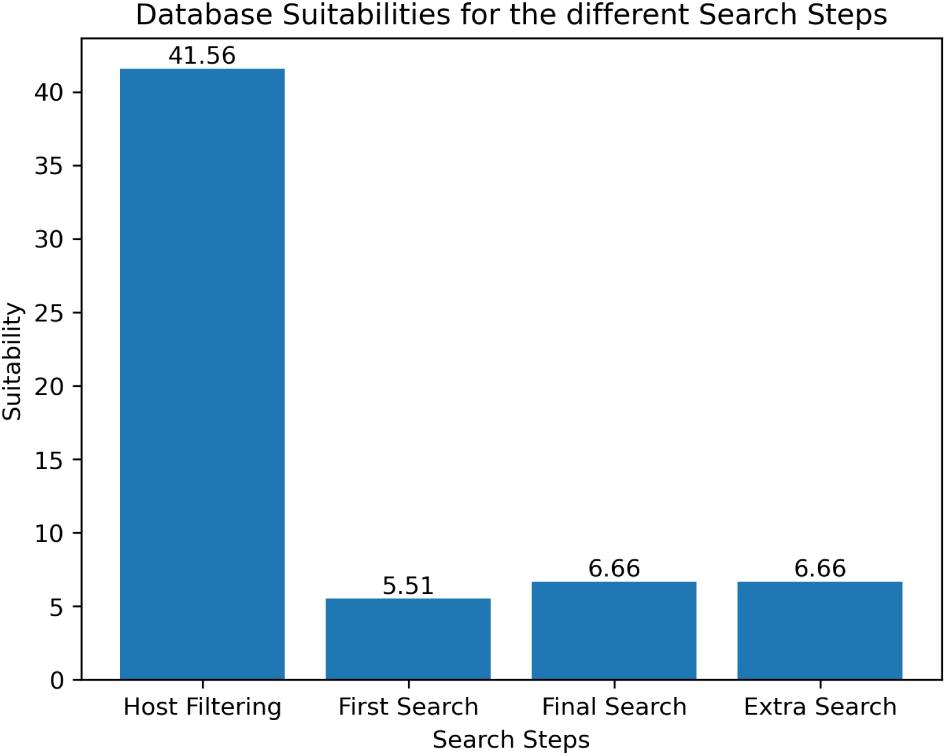
Bar plot showing the database suitabilities of the hendra sample using the query approach without the taxonomy.

To determine the threshold, for the database suitabilty, from which the user is advised to investigate the suitability of the database and the quality of the sample, the herpes sample was tested with three other, several animal host databases (*Rattus Norvegicus*, which is the actual host, *Pteropus alecto* and *Gallus gallus*). The database suitabilities dropped from 49.0% to 39.56%, 31.47% and 33.33% respectively as shown in Figure 11 in the supporting information. Therefore, it is recommended to investigate the quality of the sample as well as the suitability of the database, if the suitability is lower than 40%.

## Discussion

The results show that MultiStageSearch is able to consistently identify the correct strain of viral proteomic samples while being independent of the NCBI taxonomy. Due to the genomic data used for the strain level search, more reliable data can be used, as genomics is more established than proteomics for viral identification.^6^ Additionally, the clever generation of the proteogenomic reference database, such as filtering for duplicate proteomes, as well as the clustering of identical ORFs, helps to avoid over-representation of model strains and to improve the taxonomic identification on strain level. One example is strain “17” for the herpesvirus.

On the other hand, querying the NCBI nucleotide database using the NCBI taxonomy creates a new problem connected to the annotation of the taxonomy itself, as shown for the adenovirus sample, where “Adenoviurs 2” does not have a defined rank causing the query to use name combinations that do not find the correct strain. The major drawback of the nucleotide database is the missing annotation of strain-level Taxon-IDs for the genomes. Using the second approach of MultiStageSearch for querying the nucleotide database without the NCBI taxonomy, only using the species Taxon-ID, turns this missing annotation into an advantage by being able to find strains that are not present in the taxonomy.

There still remain more challenging species. One example is the avian bronchitis sample. Similarly to PepGM, the detected peptidomes of the top-scoring taxa are identical, even when using our proteogenomic approach. One possible explanation is that the strains “Beaudette” and “Beaudette CK” are to similar to differentiate them using mass spectrometry data. Another possibility is that not enough viral proteins are present in the sample. Figure 10 in the supporting information supports this hypothesis as even the database suitability for the genomic reference search is low with only 48.66% compared to most other samples in the benchmark, which have database suitabilities over 90% for the genomic reference search. SARS-CoV-2 is another viral species difficult to identify at strain level: due to the recent pandemic, millions of of people got infected and the virus was able to mutate at a rapid rate.^50,51^ Scientist around the world analyzed the virus and uploaded millions of uncurated proteome and genome sequences. Furthermore, there exists no subspecies or strain level classification in the NCBI taxonomy for this virus. As the correct classification of viral samples becomes even more interesting in events such as epidemic outbreaks and rapidly evolving sequence mutations, this highlights the need for methods able to identify viral strains not yet catalogued in the standard taxonomic references.

**Figure 10:**
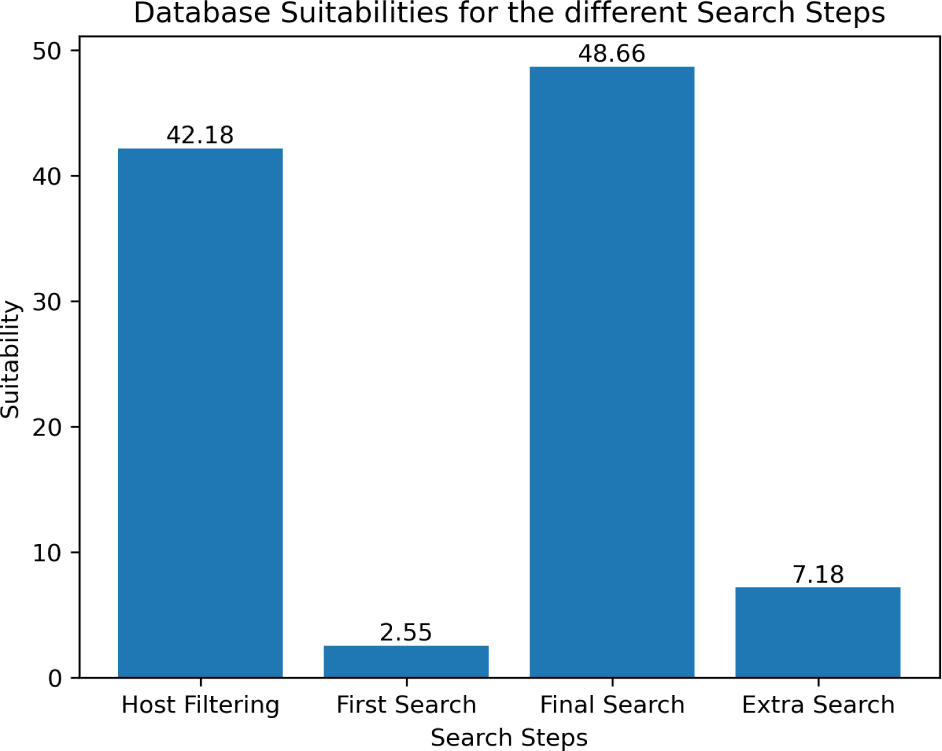
Bar plot showing the database suitabilities of the avian bronchitis sample using the query approach without the taxonomy.

**Figure 11:**
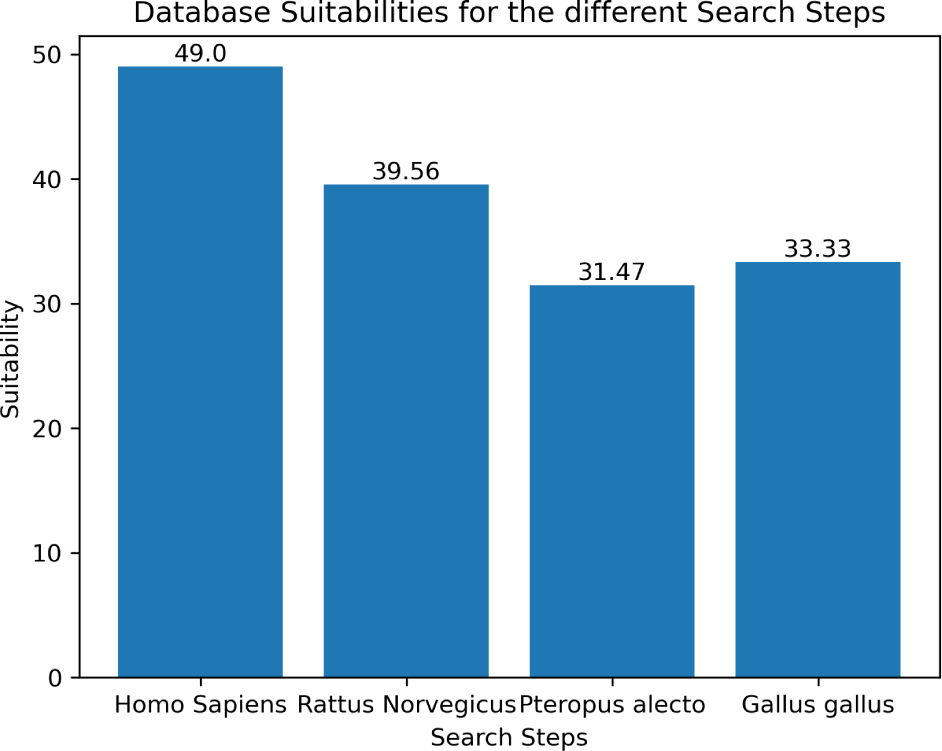
Bar plot showing the database suitabilities of the herpes sample using different host databases.

Currently, MultiStageSearch was only benchmarked on viral samples. The possibility to identify bacterial samples on strain level using MultiStageSearch is being evaluated. Bacterial samples pose additional challenges. Typically, the genomes are longer and more genomes can be found, using the query approaches of MultiStageSearch, resulting in a larger genomic reference database.

## Outlook

At the time of writing, a specialized SARS-CoV-2 mode is under developement with the goal of efficiently querying the millions of SARS-Cov-2 genomes present in the NCBI Database as well as analysing the sample to identify the lineage of the SARS-CoV-2 virus detected . Additionally, an automatically created report of the results is planned to provide the user with helpful information such as warnings for the database suitability as well as a structured overview of the results.

## Conclusion

While strain level identification using proteomics remains difficult, MultiStageSearch leverages advantages of different approaches, like proteogenomics and the novel combination of database queries and filterings. Nevertheless, there are still possibilities to improve this tool, like a specialized approach for SARS-CoV-2 samples. MutliStageSearch is implemented as a flexible snakemake workflow that easily allows the integration of additional steps and can therefore be smoothly expanded in the future to fit more individualized needs.

## Supporting Information Available

